# Marginal zone and follicular B cells respond differently to TLR4 and TLR9 stimulation

**DOI:** 10.1101/2025.05.20.655194

**Authors:** Himani Makkar, Sophie Remick, Sukanya Roy, Thomas A. Zangle, Koushik Roy

## Abstract

Marginal zone (MZ) B cells are considered to be innate-like immune cells, and follicular (FO) B cells are considered to be adaptive immune B cells. Yet, the proliferative response of MZ and FO B cells to different innate stimuli remains unclear. Here, we investigated cell growth, division, and death to determine the collective proliferative response of MZ and FO B cells in response to innate stimuli, LPS and CpG. We show that the growth rate of FO B cells is higher than that of MZ B cells in response to CpG, though MZ B cells acquire a higher mass at division, whereas both the growth rate and mass at division for MZ and FO B cells remain similar in response to LPS stimulation. We show that MZ B cells divide faster and induce a higher cRel expression in response to both CpG and LPS stimulation than FO B cells. A higher proportion of MZ B cells enter first division in response to LPS stimulation, not in response to CpG stimulation, than FO B cells. Interestingly, CpG stimulation, not LPS stimulation, leads to higher cell death in MZ B cells than FO B cells. In response to LPS stimulation, MZ B cells show a higher cell number at early time and a reduced/ similar cell number at late time, whereas in response to CpG stimulation, MZ B cells show a lower cell number at both early and late time. Our study suggests that LPS and CpG stimulation impact cell growth, division, and death differently, which in turn regulate different proliferative responses of MZ and FO B cells. Thus, our study offers a new perspective that different innate stimuli regulate different features of proliferative responses.

## Introduction

The B cell response to invading pathogens is regulated by both a non-specific response, mediated by innate immune components, and a highly specific response, mediated by adaptive immune components (1). Activation of a germline-encoded pattern-recognition receptor, such as Toll-like receptor (TLR), is a component of the innate immune response. This response serves to induce a B cell response upon recognition of pathogen-associated molecular patterns. Accordingly, both lipopolysaccharide (LPS) and CpG induce B cell activation and proliferation. LPS is a component of the outer membrane of gram-negative bacteria and a ligand for the cell surface receptor TLR4 (2). The unmethylated CpG motif-containing DNA generated from pathogens activates TLR9, an intracellular, endosomal membrane-bound receptor (2). Phenotypic, spatial, and functional analysis reveal two major subsets of mature splenic B cells: follicular (FO) and marginal zone (MZ) B cells (3). However, the regulation of cell growth and proliferation of MZ and FO B cells following innate stimulations remain unclear. Here, we investigated the cell growth and proliferation of MZ and FO B cells in response to cell surface (TLR4) and intracellular (TLR9) TLRs signals. MZ B cells are considered “innate-like” cells and exhibit higher expression of TLR4 and TLR9 compared to FO B cells (1, 4, 5). Accordingly, we expected both LPS and CpG stimulation to induce higher cell growth and proliferation in MZ B cells compared to FO B cells. Surprisingly, we revealed that LPS and CpG induce differential cell growth, proliferation, and death in MZ and FO B cells.

## Results and Discussion

### Differential growth of marginal zone and follicular B cells with TLR4 and TLR9 stimulation

Stimulation of B cells with mitogen results in cell growth due to the accumulation of cell mass and a resulting increase in cell size (6-8). Cell growth and cell division are two functionally separate processes; however, they are tightly linked and crucial for generating progenitor cells (6, 9). B cells that enter the cell growth program are protected from death and undergo cell division (8). Here, we investigated the growth of MZ and FO B cells by measuring relative cell size using forward scatter from flow cytometry (10) and cell mass using live cell quantitative phase imaging (QPI) (11-13). We found that MZ B cells are larger than FO B cells at steady state (0 hour; before stimulation) and 24 hours of stimulation with CpG and LPS (Fig. S1A-C). This observation aligns with previous studies, which show that MZ B cells are larger than FO B cells at steady state (14).

Following mitogenic stimulation, the increase in cell mass is driven by enhanced de novo synthesis of RNA, protein, and DNA, and is regulated by both anabolic and catabolic pathways. (15). We stimulated MZ and FO B cells with CpG and LPS to monitor changes in cell mass until the first division using live cell QPI (Fig. 1A). We find that MZ and FO B cells follow two trajectories of cell growth: cells that grow rapidly follow an exponential growth trajectory (Fig. 1B-E), while non-growing cells exhibit a linear growth trajectory with minimal to no growth (Fig. S1D-E). Previous studies have shown that non-growing cells undergo cell death, whereas growing cells are protected from death and enter the cell division program (7, 8). Hence, we focused on exponentially growing cells (Fig. 1B-E). We found that LPS-stimulated MZ B cells have a higher mass than FO B cells at 16 hours (MZ and FO mass: 44 pg and 39 pg, respectively) and 24 hours (MZ and FO mass: 59 pg and 50 pg, respectively) stimulation (Fig. 1F-H). Similarly, CpG-stimulated MZ B cells have a higher mass than FO B cells at 16 hours (MZ and FO mass: 48 pg and 36 pg, respectively) and 24 hours (MZ and FO mass: 64 pg and 54 pg, respectively) stimulation (Fig. 1G-I). Live cell imaging reveals that cells acquire different sizes before the first division (7). Similarly, we found that the MZ and FO B cells’ mass before the first division is widely distributed (Fig. 1J-K). We are interested in understanding whether the mass of MZ and FO cells before the first division is stimulus-dependent. Interestingly, we found that LPS-stimulated MZ and FO B cells acquire similar mass (MZ and FO mass: 66 pg and 67 pg, respectively) before first division; however, CpG-stimulated MZ B cells acquire a higher mass (83 pg) than FO B cells (74 pg), suggesting that the mass of MZ and FO before first division is stimulus-dependent (Fig. 1J-K). A previous study has shown that the cell size of founder cells before the first division determines the maximum division cycle (7). It is, therefore, possible that the mass of MZ and FO B cells before the first division could determine the maximum number of divisions, or proliferative capacity, of founder cells. It remains to be determined whether a similar mass of MZ and FO B cells before first division upon LPS stimulation results in similar division numbers, and whether a higher mass of MZ B cells before first division upon CpG stimulation leads to a higher division number.

**Figure 1:**
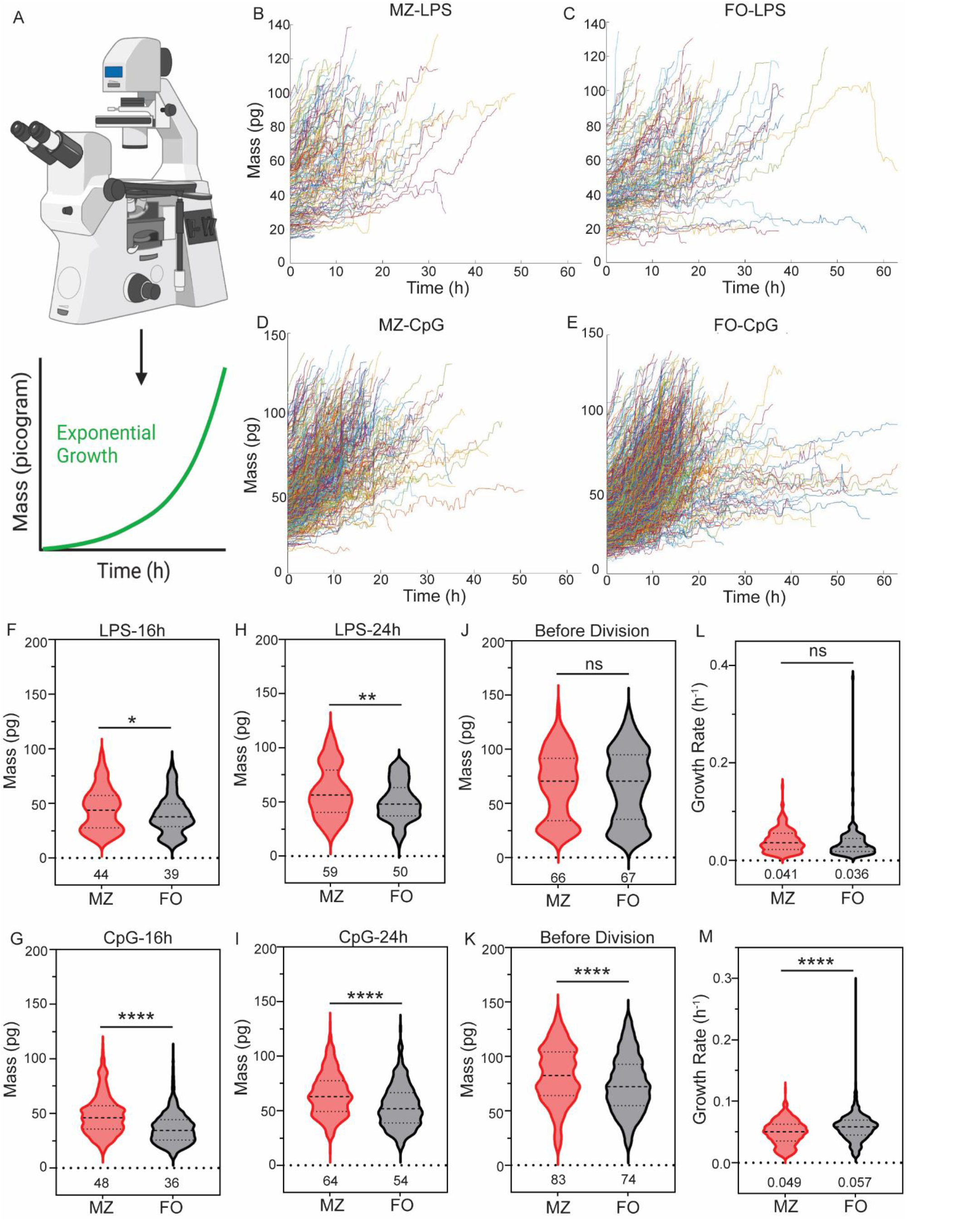
Differential growth of MZ and FO B cells. **A)** Schematic of time-lapse quantitative phase imaging to measure cell mass. Mass accumulation follows exponential growth for dividing cell. **B) + C)** Cell growth trajectories of LPS-stimulated MZ and FO B cells follow an exponential growth. The X-axis represents time in hours, and the Y-axis represents cell mass in picograms (pg). **D) + E)** Cell growth trajectories of CpG-stimulated MZ and FO B cells follow an exponential growth. **F) + H)** Violin plot of LPS-stimulated MZ and FO B cells mass at 16 h and 24 h. Cell mass is expressed in pg. The MZ and FO B cells are represented by red and black violin plots, respectively. **G) + I)** Violin plot of CpG-stimulated MZ and FO B cells mass at 16 h and 24 h. **J)** Violin plot of LPS-stimulated MZ and FO B cells mass at division. **K)** Violin plot of CpG-stimulated MZ and FO B cells mass at division. L) Violin plot of LPS-stimulated MZ and FO B cells growth rate (pg per h). **K)** Violin plot of CpG-stimulated MZ and FO B cells growth rate (pg per h). In all plots, the mean is indicated at the bottom of each group. *p < 0.05, **p < 0.01, ***p < 0.001, and not significant, ns (unpaired Student’s t test).

The joint regulation of growth and division rates determines the regulation of cell size. For example, the growth rate of mammalian cells may depend on the size of the cells immediately after birth, e.g., larger cells have a higher growth rate, and smaller cells have a lower growth rate (16). The steady state size of MZ B cells is bigger than FO B cells (Fig. S1). To determine whether the growth rate of MZ B cells is faster than that of FO B cells, we measured the mass accumulation rate, which we define as the growth rate. We found that LPS-stimulated MZ B cells have a similar growth rate (0.041 pg/h) to FO B cells (0.036 pg/h) (Fig. 1L). However, CpG-stimulated MZ B cells have a lower growth rate (0.049 pg/h) than FO B cells (0.057 pg/h) (Fig. 1M). Our date suggest that B cell growth rate depends on the type of stimulus rather than the initial size of cells. The differential regulation of growth rate in response to CpG and LPS stimulation is likely mediated by distinct signaling pathways and gene regulatory networks activated by CpG and LPS.

### Faster division of marginal zone B cells with TLR4 and TLR9 stimulation correlates with cRel expression

MZ B cells are considered “innate-like” cells that rapidly respond to stimulation and generate plasma cells independently of cell division, in contrast to FO B cells (1, 4). The proliferative properties of MZ B cells are less well understood. We labeled MZ and FO cells with the division tracking dye Cell Trace Far Red (CTFR) to study the proliferation of MZ and FO B cells, as indicated by halved CTFR intensity in each division cycle. We measured CTFR intensity at 24 and 48 hours. The CTFR plot at 24 hours shows a single peak and similar CTFR intensity for MZ and FO B cells, regardless of LPS or CpG stimulation, suggesting either no cell division or undetectable cell division at this time point (Fig. 2A and 2E). Interestingly, CTFR intensity is dramatically reduced in MZ B cells compared to FO B cells at 48 hours following both LPS and CpG stimulation, suggesting that MZ B cells divide more quickly than FO B cells in response to innate stimulation (Fig. 2B and 2F). To quantify first division time (Tdiv0), we utilized the phenotyping computational tool FlowMax (17, 18). FlowMax uses a modified cyton model to estimate Tdiv0 from the dye dilution time course. Upon LPS stimulation, the estimated Tdiv0 is 32.7 hours for MZ B cells and 42.4 hours for FO B cells (Fig. 2C and 2D). Upon CpG stimulation, the estimated Tdiv0 is 26.3 hours for MZ B cells and 37.5 hours for FO B cells (Fig. 2G and 2H). Thus, our study demonstrated that MZ B cells divide more rapidly than FO B cells in response to both CpG and LPS stimulation, suggesting that TLR stimulation prompts a faster proliferative response in MZ B cells. Hence, MZ B cells possess a unique property that enables them to divide and differentiate into antibody-producing plasma cells more rapidly compared to FO B cells. MZ B cells are predominantly located in the marginal zone at the white and red pulp interface, and FO B cells are predominantly located in the follicle in the spleen (1). The location of MZ B cells in the spleen enables them to encounter bloodborne pathogens and circulating stimuli earlier than FO B cells, resulting in faster stimulation (1). A previous study shows that T-cell independent antigen-specific B cell response is predominantly from MZ B cells upon immunization with bacteria (19). Faster MZ B cell division enables the rapid expansion of antigen-specific B cell clones.

**Figure 2:**
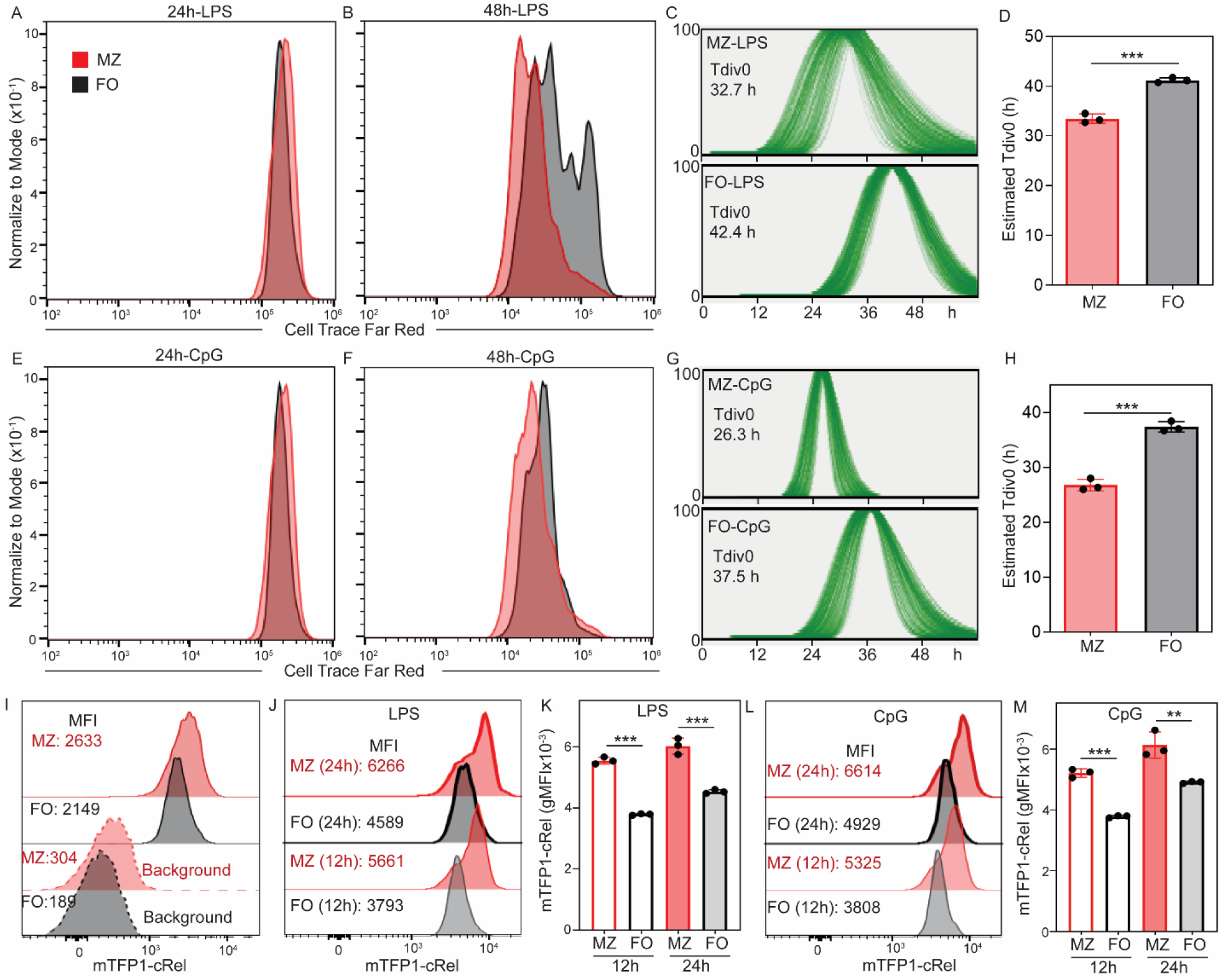
Faster division of MZ B cells is correlated with cRel expression. **A) + B)** Cell Trace Far Red (CTFR) histogram of MZ and FO B cells at 24 h and 48 h following stimulation with LPS. Data shown are representative of 3 biological replicates. The X-axis represents the dilution of CTFR, and the Y-axis represents the histogram, normalized to the mode. **C)** Distribution of first division time (Tdiv0) of MZ (upper panel) and FO (upper panel) B cells stimulated with LPS were analyzed using FlowMax, running Fcyton model to predict Tdiv0. Data shown are representative of 3 biological replicates. **D)** Bar graph shows the estimated Tdiv0 of MZ and FO B cells stimulated with LPS. **E) + F)** CTFR histogram of MZ and FO B cells at 72 h and 96 h following stimulation with CpG. Data shown are representative of 3 biological replicates. The X-axis is the dilution of CTFR, and the Y-axis is the histogram normalized to mode. **G)** Distribution Tdiv0 of MZ (upper panel) and FO (upper panel) B cells stimulated with CpG were analyzed using FlowMax, running Fcyton model to predict Tdiv0. Data shown are representative of 3 biological replicates. **H)** Bar graph shows the estimated Tdiv0 of MZ and FO B cells stimulated with CpG. **I)** Total cellular cRel fluorescence histogram between MZ (dark red) and FO (dark black) B cells without stimulation (0 h). Background fluorescence of MZ and FO B cells are shown by light red and light black color. For background fluorescence, MZ and FO B cells were isolated C57BL/6 mice expressing cRel without mTFP1 knock-in. Data shown are representative of 3 biological replicates. **J) + K)** Total cellular cRel fluorescence histogram between MZ (dark red) and FO (dark black) B cells stimulated with LPS for 14 h and 24 h. **L) + M)** Total cellular cRel fluorescence histogram between MZ (dark red) and FO (dark black) B cells stimulated with CpG for 14 h and 24 h. The number of dots in each plot is the number of replicates. In all plots, the mean is indicated at the bottom of each group. *p < 0.05, **p < 0.01, ***p < 0.001, and not significant, ns (unpaired Student’s t test).

Therefore, combining faster division and faster plasma cell generation enables MZ B cells to enrich an early antigen-specific B cell response, providing the first line of antibody-mediated defense against bacterial infection.

Previous studies have demonstrated that stimulating naïve B cells with CpG or LPS activates the NFκB signaling pathway (20-22). Furthermore, deletion of cRel, a key NFκB subunit, impairs B cell proliferation and survival (8, 21, 23). Partial deletion of cRel in B cells leads to reduced DNA synthesis, indicating impaired cell cycle progression. Although transgenic expression of the anti-apoptotic factor Bcl2 in cRel-deficient B cells prevents cell death, it does not restore cell division. These findings suggest that cRel is specifically required for B cell cycle progression, independent of its role in cell survival (6, 21, 23). In line with this, our recent study shows that high cRel-expressing FO B cells induced a higher cRel expression and divided faster than low cRel-expressing FO B cells, and, further, we show that cRel expression is higher in MZ B cells compared to FO B cells (Fig. 2I) (24). To test whether the faster division of MZ B cells correlates with higher induced cRel expression before the first division, we measured the kinetics of cRel expression by flow cytometry using cRel reporter (mTFP1-cRel) B cells at 12 and 24 hours. Upon LPS stimulation, the geometric mean fluorescence intensity (gMFI) at 12 hours is 5661 and 3737 for MZ and FO B cells, respectively, and the gMFI at 24 hours is 6266 and 4589 for MZ and FO B cells, respectively (Fig. 2J-K). Upon CpG stimulation, gMFI at 12 hours is 5325 and 3808 for MZ and FO B cells, respectively, and gMFI at 24 hours is 6614 and 4929 for MZ and FO B cells, respectively (Fig. 2L-M). We showed that MZ B cells induced a higher cRel expression than FO B cells before first division (12- and 24-hours) with both CpG and LPS stimulation (Fig. 2J-M). Thus, faster division of MZ B cells is correlated with both higher steady-state and induced cRel expression in MZ B cells.

### Differential proliferation of MZ and FO B cells in response to TLR4 and TLR9 stimulation

Previous studies have shown that LPS stimulation of MZ B cells induces higher proliferation than FO B cells (5, 14), and surprisingly, CpG stimulation induces similar proliferation of MZ and FO B cells (1, 4, 5). However, the proliferation kinetics and the proportion of cells entering the proliferative program are understudied. We measured the kinetics of proliferation using the CTFR dilution and by counting the number of cells. We found that both MZ and FO B cells reach a similar maximum cell number at 72 hours upon LPS stimulation, and FO B cells may have a higher cell number at 96 hours than MZ B cells (Fig. 3A-C). Interestingly, MZ B cells have a higher cell number at 48 hours (Fig. 3C). The increase in cell number for MZ B cells at 48 hours may be due to the following reasons: (1) faster cell division, (2) lower cell death, and (3) a higher proportion of cells entering cell division. We found that MZ B cells divide faster than FO B cells in response to LPS stimulation (Fig. 2D). We then measured cell death and found that a similar proportion of MZ and FO B cells undergo death at 48 hours (Fig. 3D-F). We estimated the proportion of cells entering the first division using FlowMax and found that a higher proportion of MZ B cells enter the first division than FO B cells upon LPS stimulation (Fig. 3G). Our study suggests that a higher MZ B cell number at 48 hours of LPS stimulation combines faster division and a higher proportion of cells entering the first division cycle. We found that FO B cells have a higher cell number and proliferation than MZ B cells at 72 and 96 hours of CpG stimulation (Fig. 3H-J). Interestingly, we found that CpG stimulation induces higher cell death in MZ B cells (Fig. 3K-M) and faster division compared to FO B cells at 48 h (Fig. 2H). FlowMax analysis showed that a similar proportion of MZ and FO B cells enter the first division cycle (Fig. 3N). Therefore, our study suggests that the higher number of FO B cells in response to CpG stimulation is due to reduced cell death of FO B cells. Overall, we demonstrated that LPS and CpG stimulation induce faster division in MZ B cells. However, LPS and CpG stimulation also induce differential cell death and proliferation in MZ and B cells, suggesting stimulus-dependent modulation of cell death and proliferative programs.

**Figure 3:**
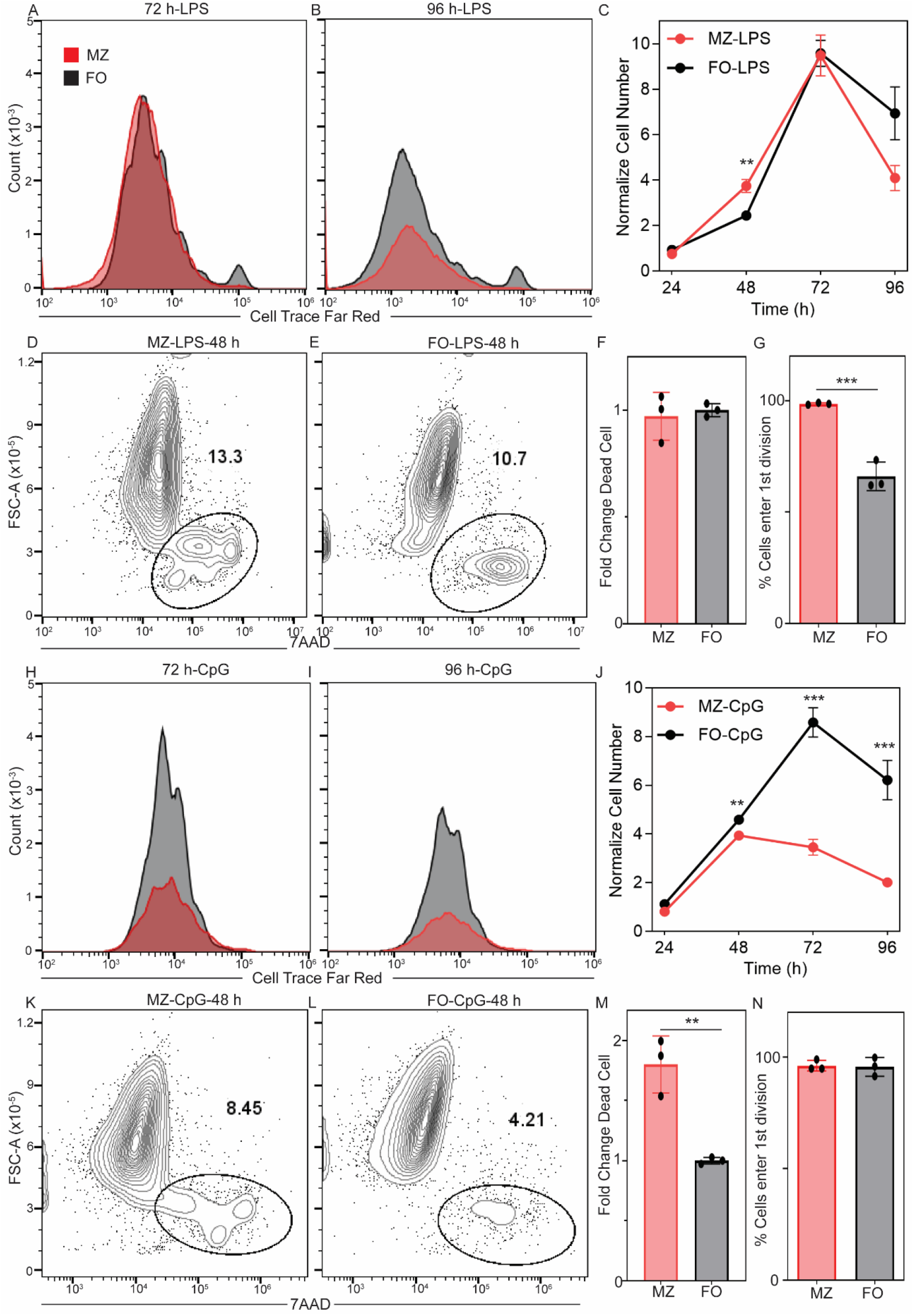
Sustained proliferation of FO B cells upon TLR4 and TLR9 stimulation. **A) + B)** CTFR histogram of MZ and FO B cells at 72 h and 96 h following stimulation with LPS. The X-axis is the dilution of CTFR, and the Y-axis is cell counts. **C)** Normalize cell number of MZ and FO B cells at 24 h, 48 h, 72 h and 96 h of three replicates. Normalized cell number is calculated by the cell number at a particular time/ seeding cell number at 0 h. The X-axis is time in hours, and the Y-axis is normalized cell number. **D) + E)** Dead cells proportion of MZ and FO B cells following stimulation with LPS for 48 h. The X-axis is 7AAD, and the Y-axis is forward scattered area. Data shown are representative of 3 biological replicates. **F)** Bar graph shows fold change of dead cell between MZ and FO B cells stimulated with LPS for 48 h. Fold change is calculated by % of dead cells in MZ/ FO. **G)** Bar graph shows the predicted proportion of cells entering 1^st^ division cycle in MZ and FO B cells stimulated with LPS and data were analyzed using FlowMax, running Fcyton model. **H) + I)** CTFR histogram of MZ and FO B cells at 72 h and 96 h following stimulation with CpG. The X-axis is the dilution of CTFR, and the Y-axis is cell counts. **J)** Normalize cell number of MZ and FO B cells at 24 h, 48 h, 72 h and 96 h of three replicates. Normalize cell number is calculated by the cell number at a particular time/ seeding cell number at 0 h. The X-axis is time in hours, and the Y-axis is normalized cell number. **K) + L)** Dead cells proportion of MZ and FO B cells following stimulation with CpG for 48 h. The X-axis is 7AAD, and the Y-axis is forward scattered area. Data shown are representative of 3 biological replicates. **F)** Bar graph shows fold change of dead cells between MZ and FO B cells stimulated with CpG for 48 h. Fold change is calculated by % of dead cells in MZ/ FO. **G)** Bar graph shows the predicted proportion of cells entering 1^st^ division cycle in MZ and FO B cells stimulated with CpG and data were analyzed using FlowMax, running Fcyton model. The number of dots in each plot is the number of replicates. In all plots, the mean is indicated at the bottom of each group. *p < 0.05, **p < 0.01, ***p < 0.001, and not significant, ns (unpaired Student’s t test).

In this work, we investigated whether MZ and FO B cells differ in cell growth and proliferative properties, and whether these properties are dependent on the type of stimulus. We found that CpG-stimulated FO B cells have a higher growth rate compared to MZ B cells, whereas LPS-stimulated FO and MZ B cells have a similar growth rate, suggesting stimulus-dependent regulation of growth rate for MZ and FO B cells (Fig. 1L and 1M). MZ B cells divide more rapidly than FO B cells in response to both CpG and LPS stimulation (Fig. 2D and 2H). Interestingly, the proportion of cells that enter the first division is higher for MZ B cells compared to FO B cells in response to LPS stimulation, whereas the proportion remains similar for MZ and FO B cells in response to CpG stimulation (Fig. 3G and 3N). Thus, our study suggests that the growth rate and proliferative properties of MZ and FO B cells are stimulus-dependent. The differential regulation of growth and proliferative properties of MZ and FO B cells is likely linked with signaling pathways and gene regulatory networks activated by CpG and LPS stimulation, and remains to be determined.

## Methods

### Marginal zone and follicular B cell isolation and proliferation assay

Spleens were harvested and homogenized from 10-14 week old C57BL/6 and mTFP1-cRel mice. Marginal zone and follicular B cells were isolated using magnetic microbeads as recommended by the manufacturer’s protocol (Miltenyi Biotec #130-100-366) and described previously (17). Briefly, spleenocytes were incubated with the marginal zone and follicular B cell biotin-antibody cocktail in MACS buffer (Phosphate buffer saline, pH 7.4, 0.5% Bovine serum albumin and 2 mM Ethylenediaminetetraacetic acid, pH 8) for 5 minutes at 4-8°C, followed by incubation with streptavidin magnetic beads for 10 minutes at 4-8°C. Cells were washed with MACS buffer and passed through the LS column (Miltenyi Biotec #130-042-401) to collect total B cells in the flowthrough. Collected total B cells were incubated with anti-CD23 magnetic beads for 15 minutes at 4-8°C, washed with MACS buffer, and passed through the LS column to enrich MZ and FO B cells.

MZ and FO B cells were labeled with 1 μM Cell Trace Far Red (CTFR) (Thermo Fisher Scientific #C34572), and proliferation was measured as described previously (25). CTFR labelled MZ and FO stimulated with 250 nM of CpG ODN 1668 (5’-tccatgacgttcctgatgct-3’ synthesized at the University of Utah core facility) and 10 μg/mL of LPS from *Escherichia coli* O55:B5 (Sigma-Aldrich #L6529) at 50,000 cells in 250 uL freshly prepared B cell media (1640 RPMI, 10% FBS, 1X pen-strep, 5 mM glutamine, 1 mM sodium pyruvate, 1 mM MEM non-essential amino acid, 20 mM HEPES and 55 μM 2-mercaptoethanol) in a 48-well plate 37°C, 5% CO_2_ containing humidified chamber for a period of 4 days. B cell proliferation and death were measured at 24-, 48-, 72-, and 96-hours using flow cytometry (Beckman Coulter Cytoflex). Cell death was determined by staining the cells with 7-AAD (Biolegend #420404).

### Cell biological parameter estimation (FlowMax)

CpG (250 nM) and LPS (10 μg/mL) stimulated CTFR labelled MZ and FO B cells were acquired at 14-, 24-, 48-, 72-, and 96-h using flow cytometry (Beckman Coulter Cytoflex), and the volume of acquisition was 30-100 uL. Live cells were gated, and cell acquisition volume was normalized on the FlowMax software as described earlier (17, 18). Briefly, The CTFR fluorescence of the generation 0 cells (undivided peak) was identified manually for each time point, and the fcyton model was used to estimate the cell biological parameters of first division time (Tdiv0) and the proportion of cells that enter 1^st^ division cycle. We performed at least 500 simulations to estimate the maximum likelihood of cell biological parameters.

### Quantitative phase imaging (QPI)

QPI was performed on a custom built (13, 26), differential phase contrast microscope consisting of an LED array (Adafruit) sequenced by a microcontroller (Arduino), a 10X NA = 0.25 objective (Olympus), a f = 200 mm tube lens (Thorlabs) and a 1920×1200 megapixel Grasshopper 3 CMOS camera (FLIR). Positioning was controlled by a programmable *xy* stage (Thorlabs) and a *z* stage (Thorlabs) driven by a stepper motor (Mouser). Quantitative phase images were reconstructed using Tikhonov regularization (27, 28). A cell average specific refractive increment of 1.8×10^−4^ m^3^/kg was used for calculation of cell mass (15, 29). Primary, naïve B cells were plated at 25,000 cells/mL in a 96 well plate (Eppendorf) coated with Poly-D-Lysine (Gibco), and stimulated 16 h before imaging to allow the plate to equilibrate before the start of imaging.

## Author Contributions

K.R. and T.Z. designed the study. H.M., S. Roy, and K.R. performed the B-cell proliferation assay and analysis. S. Remick performed the imaging experiment and analysis. K.R. wrote the manuscript. T.Z. edited the manuscript. All authors provided feedback on the writing and agreed to the final version of the manuscript.

## Acknowledgments

We are thankful to the current and former lab members of the Roy lab and the Zangle lab for critical discussion and feedback on this study. K.R. acknowledges funding from the Center of Aging, and the Department of Pathology, University of Utah, and the National Institute of Allergy and Infectious Diseases (R56AI177789). T.Z. acknowledges funding from the Department of Defense, Congressionally Directed Medical Research Program (BC230633). Figure 1 is created with Biorender.com.

**Figure S1:**
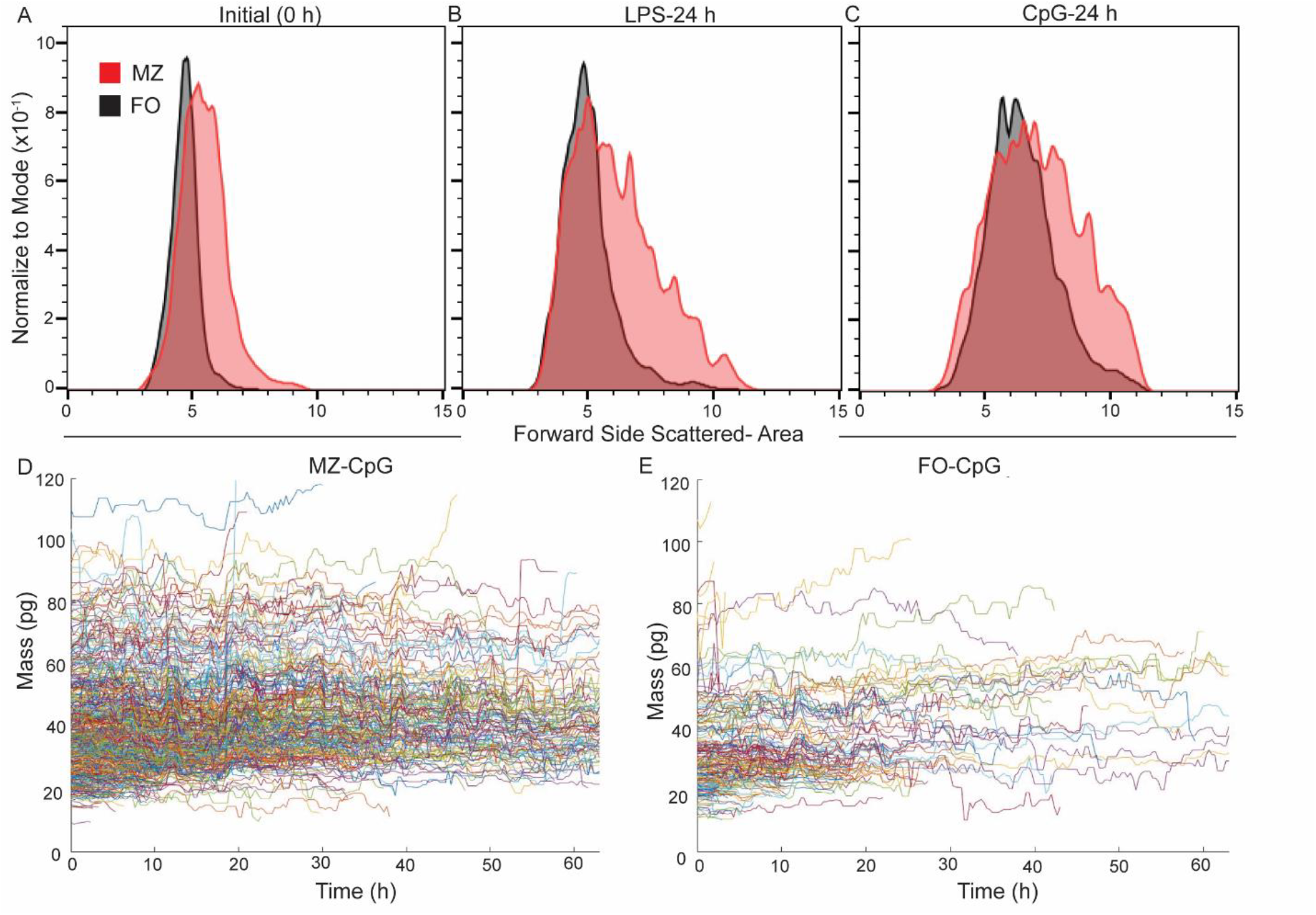
MZ B cells are larger than FO B cells. The size of MZ and FO B cells was measured using flow cytometry, as determined by forward-scattered light. **A)** size at 0 h, **B)** size at 24 h following LPS stimulation, and **C)** size at 24 h following CpG stimulation. The MZ and FO B cells are represented by red and black histograms, respectively. **D)** and **E)** Linear cell growth trajectories of CpG-stimulated MZ and FO B cells. The X-axis represents time in hours, and the Y-axis represents cell mass in picograms (pg).

